# Quantification of microaerobic growth of *Geobacter sulfurreducens*

**DOI:** 10.1101/597229

**Authors:** Christina E. A. Engel, David Vorländer, Rebekka Biedendieck, Rainer Krull, Katrin Dohnt

## Abstract

*Geobacter sulfurreducens* was originally considered a strict anaerobe. However, this bacterium was later shown to not only tolerate exposure to oxygen but also to use it as terminal electron acceptor. Research performed has so far only revealed the general ability of *G. sulfurreducens* to reduce oxygen, but the oxygen uptake rate has not been quantified yet, nor has evidence been provided as to how the bacterium achieves oxygen reduction. Therefore, microaerobic growth of *G. sulfurreducens* was investigated here with better defined operating conditions as previously performed and a transcriptome analysis was performed to elucidate possible metabolic mechanisms important for oxygen reduction in *G. sulfurreducens*. The investigations revealed that cell growth with oxygen is possible to the same extent as with fumarate if the maximum specific oxygen uptake rate (sOUR) of 95 mg_O2_ g_CDW_ ^-1^ h^-1^ is not surpassed. Hereby, the entire amount of introduced oxygen is reduced. When oxygen concentrations are too high, cell growth is completely inhibited and there is no partial oxygen consumption. Transcriptome analysis suggests a menaquinol oxidase to be the enzyme responsible for oxygen reduction. Transcriptome analysis has further revealed three different survival strategies, depending on the oxygen concentration present. When prompted with small amounts of oxygen, *G. sulfurreducens* will try to escape the microaerobic area; if oxygen concentrations are higher, cells will focus on rapid and complete oxygen reduction coupled to cell growth; and ultimately cells will form protective layers if a complete reduction becomes impossible. The results presented here have important implications for understanding how *G. sulfurreducens* survives exposure to oxygen.

## Introduction

*Geobacter sulfurreducens,* a δ-proteobacterium found in subsurface environments, was originally reported to be a strict anaerobe (1). However, it was later shown that this bacterium can grow with oxygen as terminal electron acceptor when 5 % of oxygen or less are added to the headspace of cultivation flasks (2). But this study did not investigate dissolved oxygen concentrations reached in the liquid phase under these conditions (2). Further, the analysis of the genome of *G. sulfurreducens* revealed several enzymes that could be involved in the reduction of oxygen (3). The expression of many of the corresponding genes is regulated by the RpoS regulon in *G. sulfurreducens*, which was shown to be inevitable for cell growth with oxygen (4). Part of the RpoS regulon are genes for the cytochrome c oxidase, which is thought to be most likely responsible for cell growth with oxygen, but genes for a cytochrome d ubiquinol oxidase and a rubredoxin-oxygen oxidoreductase have also been found to be regulated by RpoS and encode for enzymes that may play a role in oxygen reduction (3,5). The gene for a cytochrome d ubiquinol oxidase has also been found upregulated in *Geobacter uraniireducens*, a close relative of *G. sulfurreducens*, after exposure to oxygen (6). None of these studies have, however, compared transcriptome levels of wild type cells grown with oxygen as terminal electron acceptor to cells grown under anaerobic conditions.

Further studies have examined the response of *G. sulfurreducens* triggered by oxygen exposure, investigating for example the function of hydrogenases or genes involved in the regulation of cell growth (7,8). But the exact mechanisms of how *G. sulfurreducens* deals with oxygen are still mostly subject to speculation.

To address the described issues, this study aims to quantify the maximum specific oxygen uptake rate (sOUR) of *G. sulfurreducens*. To achieve this, cultivations with altering amounts of oxygen were performed. The cells’ reactions to altering oxygen amounts are investigated by transcriptome analysis. This analysis aims at determining the enzymes responsible for the reduction of oxygen by *G. sulfurreducens*, especially in relation to the enzymes already reported to be most likely involved in this process. Thus, the intention of the present work is to enhance the current knowledge of the microaerobic growth of *G. sulfurreducens*.

## Results

### Comparison between microaerobic and anaerobic growth

*G. sulfurreducens* was cultivated with altering oxygen concentrations and under anaerobic conditions. The corresponding growth curves are depicted in **Fig. 1**. An anaerobic cultivation with 40 mM of fumarate provided in access as terminal electron acceptor (40F0%, white squares) was performed to serve as control experiment. Cells grew exponentially in this experiment with a specific growth rate of 0.109 ± 0.015 h^-1^, reaching a maximum cell density of 0.417 ± 0.008 g_CDW_ L^-1^. In a second anaerobic experiment, the amount of fumarate was reduced to 4 mM to create electron acceptor limited conditions (4F0%, white triangles in **Fig. 1**). Cell growth was strongly reduced and no longer exponential, but rather linear. A small increase in cell dry weight could still be observed.

**Fig. 1.**
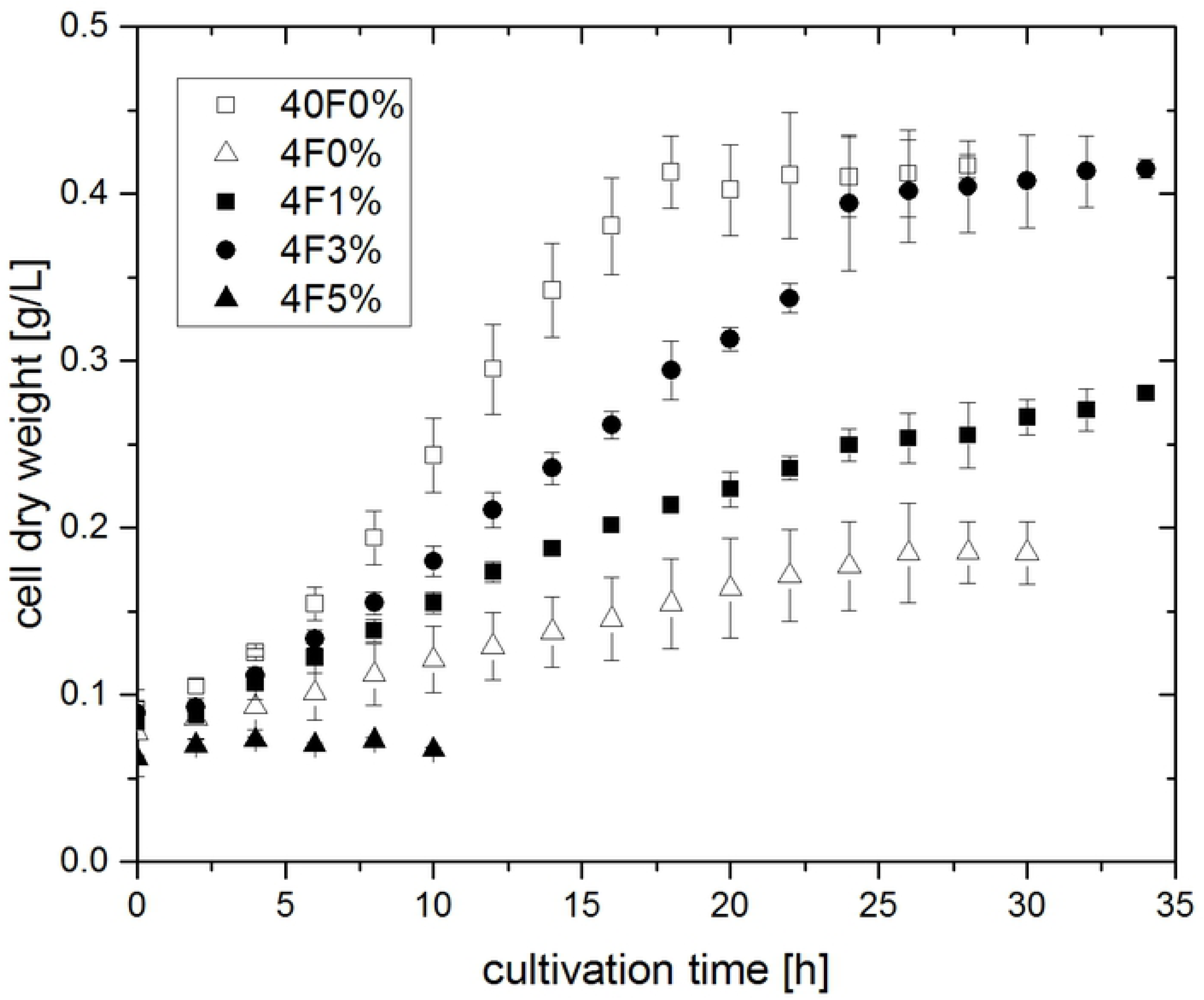
Cell growth of *G. sulfurreducens* at 30 °C with altering oxygen (between 0 and 5 %) and fumarate (4 or 40 mM) concentrations. Error bars indicate standard deviation from biological triplicates.

In the following three microaerobic cultivations, cells were provided with oxygen as an additional and alternative electron acceptor next to 4 mM of fumarate. As it was already shown by Lin et al. that *G. sulfurreducens* can only reduce oxygen after cell growth with fumarate (2), microaerobic cultivations were performed with an initial anaerobic adjustment period of 2 h. Afterwards, the gas composition was changed to contain 1, 3 or 5 % of oxygen (4F1%, 4F3% and 4F5%, black symbols in **Fig. 1**). Linear cell growth could be observed in the cultivations provided with 1 and 3 % of oxygen. Hereby biomass increase occured faster compared to experiment 4F0%. Towards the end of experiment 4F3%, there was even the same maximum biomass concentration of 0.415 ± 0.006 g_CDW_ L^-1^ achieved, as was seen in the control experiment with 40 mM of fumarate. The experiment that was performed with 5 % of oxygen in the gas inlet did not result in any cell growth.

During cultivations, the dissolved oxygen (DO) concentration within reactors was recorded and the concentration of acetate was determined by HPLC analysis. Exemplary, the biomass and acetate concentration of experiment 4F3% are plotted against the cultivation time in **Fig. 2**, together with the DO concentration. The change in gas composition after 2 h could be observed by a short spike in the DO curve. Immediately following this, the DO returned to values close to 0 mg L^-1^. A linear increase of the biomass concomitant with a linear decrease of the acetate concentration could be seen from 2 to 24 h of the cultivation. After 24 h of cultivation, cell growth and acetate consumption both subsided. Around this time, the DO concentration abruptly increased to approximately 1 mg L^-1^. As this did not happen at the exact same time point in each reactor, the average DO curve shows a very large deviation here.

**Fig. 2.**
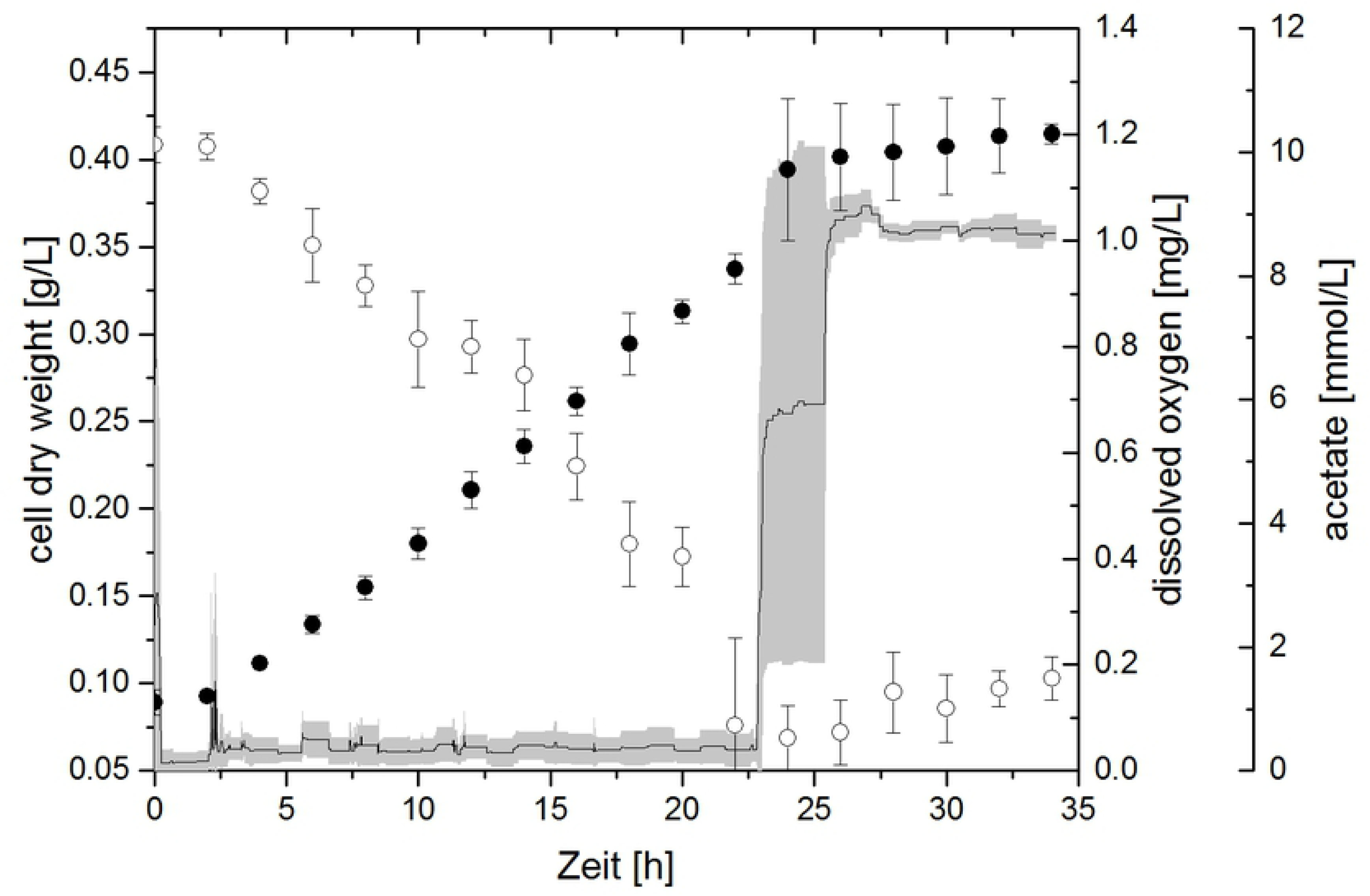
Cell dry weight (filled circles), acetate concentration (open circles) and dissolved oxygen concentration (line) during experiment 4F3% (4 mM fumarate, 3 % oxygen). Error bars (acetate and cell dry weight) and grey area (dissolved oxygen) indicate standard deviation of biological triplicates.

In the experiment 4F1% DO concentrations remained at 0 mg L^-1^ for the entire cultivation time. When 5 % of oxygen was provided, however, DO rose to 1.4 mg L^-1^. Acetate was consumed linearly in experiments 4F0% and 4F1%, but there was no significant acetate consumption in experiment 4F5%.

A final microaerobic experiment was performed in which the amount of oxygen in the gas inlet was increased stepwise dependent on the biomass concentration present (4FSI, see **Fig. 3**). Like in the other experiments, the DO concentration remained at 0 mg L^-1^ for the entire cell growth time. After cell growth stopped and acetate was depleted completely, DO increased to approximately 3.7 mg L^-1^, which is in accordance with the amount of oxygen in the gas inlet being set to 12.5 % towards the end of the growth phase. In contrast to the other microaerobic experiments, the cell growth observed was exponential. The maximum specific growth rate achieved in this way was determined to be 0.124 ± 0.007 h^-1^. Further, a maximum biomass concentration of 0.473 ± 0.018 g_CDW_ L^-1^ was achieved, which is 13 % higher than the biomass concentration seen in the anaerobic control (compare **Fig. 1**, experiment 40F0%).

**Fig. 3.**
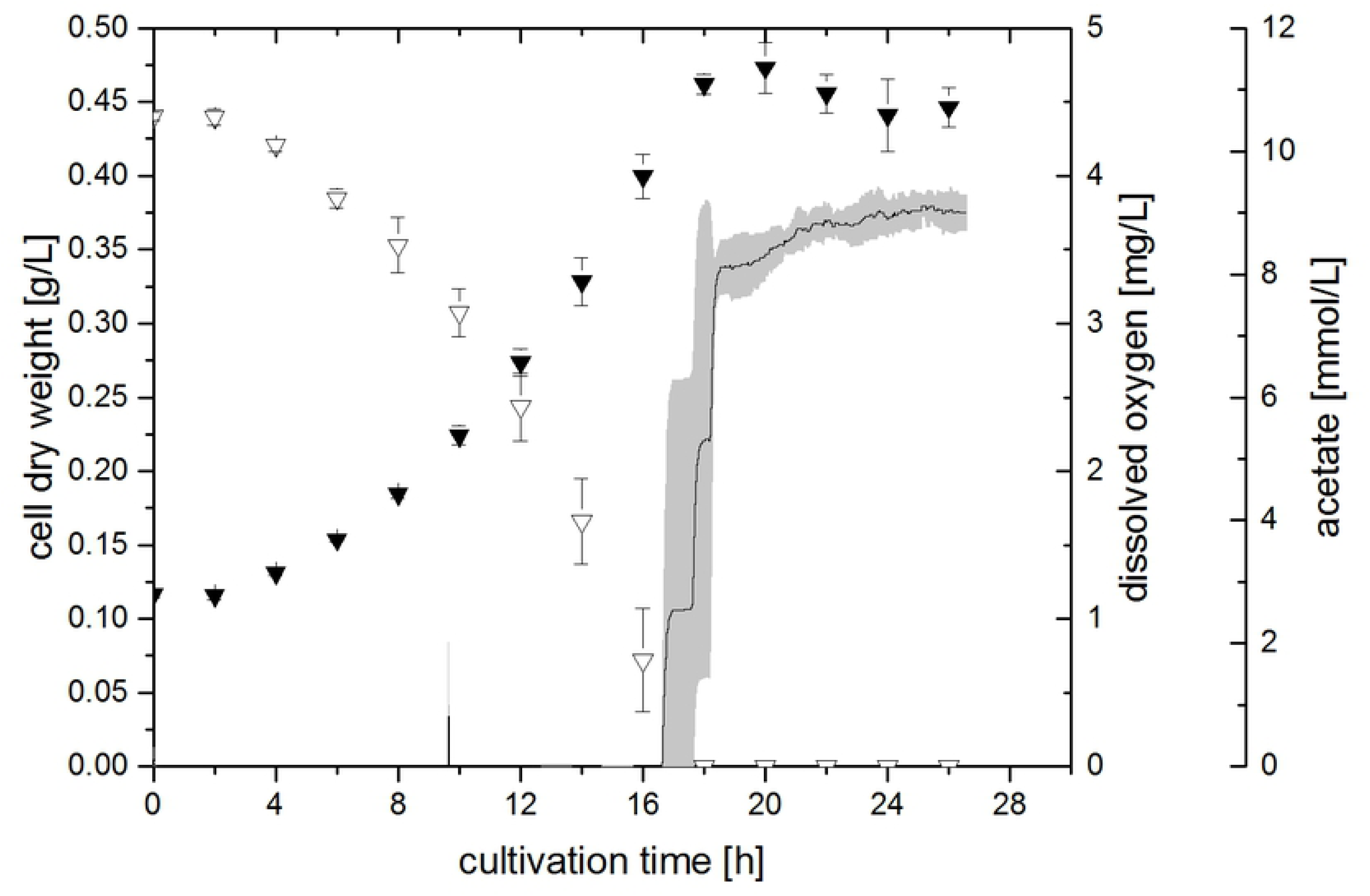
Cell dry weight (filled triangles), acetate concentration (open triangles) and dissolved oxygen concentration (line) during experiment 4FSI (4 mM fumarate, stepwise increasing amount of oxygen). Error bars (acetate and cell dry weight) and grey area (dissolved oxygen) indicate standard deviation of biological triplicates.

### Concentration profiles of succinate, fumarate and pyruvate

Apart from acetate, the concentration of succinate, fumarate, pyruvate and 2-oxoglutarate was also measured during cultivations. 2-oxoglutarate was not found in significant amounts in any experiment.

Concentration profiles of fumarate and succinate of experiments 4F0%, 4F1%, 4F3% and 4FSI are depicted in **Fig. 4**. In all experiments a consumption of fumarate and a simultaneous accumulation of succinate could be observed at first. In the case of the experiments 4F0% and 4F1%, this trend was continued until the fumarate concentration reached levels close to 0 mM (approximately after 20 h of cultivation). After that time, there was no further change in concentration levels of either substance. Contrary to this, succinate concentration was decreasing again after 24 h and 10 h in the experiments 4F3% and 4FSI, respectively. There was also a corresponding slight increase in fumarate in these experiments, although this increase was not as pronounced as the degradation of succinate.

**Fig. 4.**
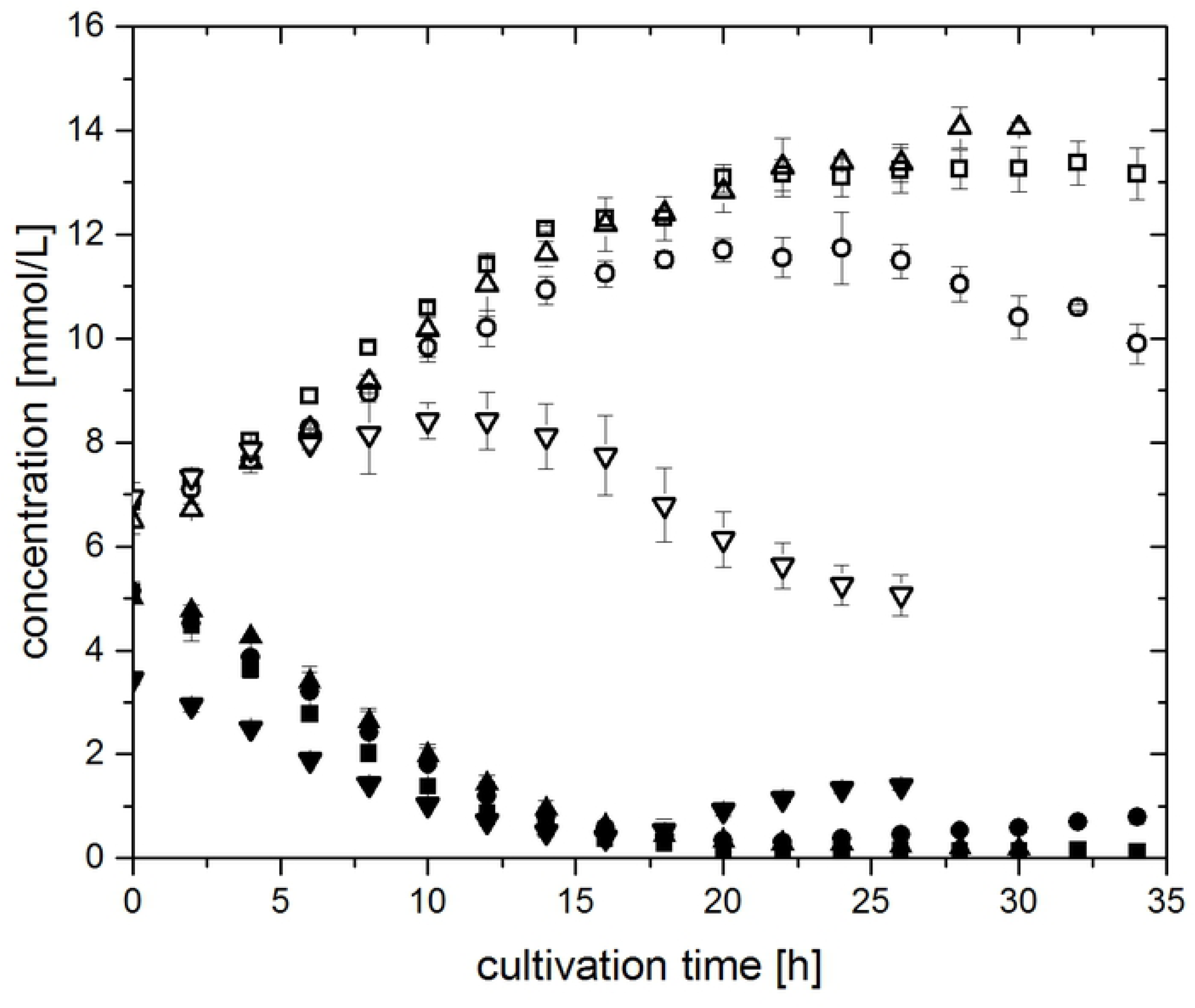
Concentration profiles of fumarate (closed symbols) and succinate (open symbols) during experiments 4F0% (4 mM fumarate, 0 % oxygen, up-pointing triangles), 4F1% (4 mM fumarate, 1 % oxygen, squares), 4F3% (4 mM fumarate, 3 % oxygen, circles) and 4FSI (4 mM fumarate, stepwise increasing oxygen amount, down-pointing triangles). Error bars indicate standard deviation of biological triplicates.

The concentration profile of pyruvate can be seen in the supplementary material in **S1_Fig** for all cultivations. Three different trends were observed. Pyruvate was present at concentrations below 0.4 mM from the preculture used for inoculation. In experiment 40F0%, pyruvate concentration increased to 3.3 mM. In cultivations 4F0% and 4F1%, pyruvate did not accumulate, but was rather degraded to 0.1 mM. However, in experiments 4F3% and 4FSI, there appeared to be an accumulation of pyruvate in the medium as long as the sOUR was high. In the case of experiment 4FSI, a final pyruvate concentration of 0.73 mM is obtained.

### Transcriptome analysis

To investigate the potential mechanisms of *G. sulfurreducens* for dealing with oxygen, a microarray transcriptome analysis of the cultivations 4F1%, 4F3% and 4F5% was performed, comparing the transcription levels of these experiments with the cultivation 40F0%. The expression of 19, 85 and 558 genes was upregulated while the expression of 1, 33 and 483 genes was downregulated, respectively (mean |logFC| ≥ 1). A selection of genes differently expressed can be found in **Table 1**. Data for all genes and experiments can be found in the supporting information in **S1_table** (for experiment 4F1%), **S2_table** (for experiment 4F3%) and **S3_table** (for experiment 4F5%). Of the genes corresponding to enzymes involved in oxygen reduction, only GSU1641 was found to be expressed at higher levels in experiments 4F1% and 4F3%. This gene encodes for the catalytic subunit of the cytochrome bd menaquinol oxidase. A number of type IV pili genes were expressed at higher levels in experiment 4F1%, but not in the others. Genes encoding for enzymes from the central carbon and energy metabolism were mainly not expressed differentially in experiment 4F1% and 4F3%, but the transcription of many of them was downregulated in experiment 4F5%. Genes corresponding for enyzmes involved in biofilm formation and encapsulation were expressed at higher levels in experiment 4F5%.

**Table 1.**
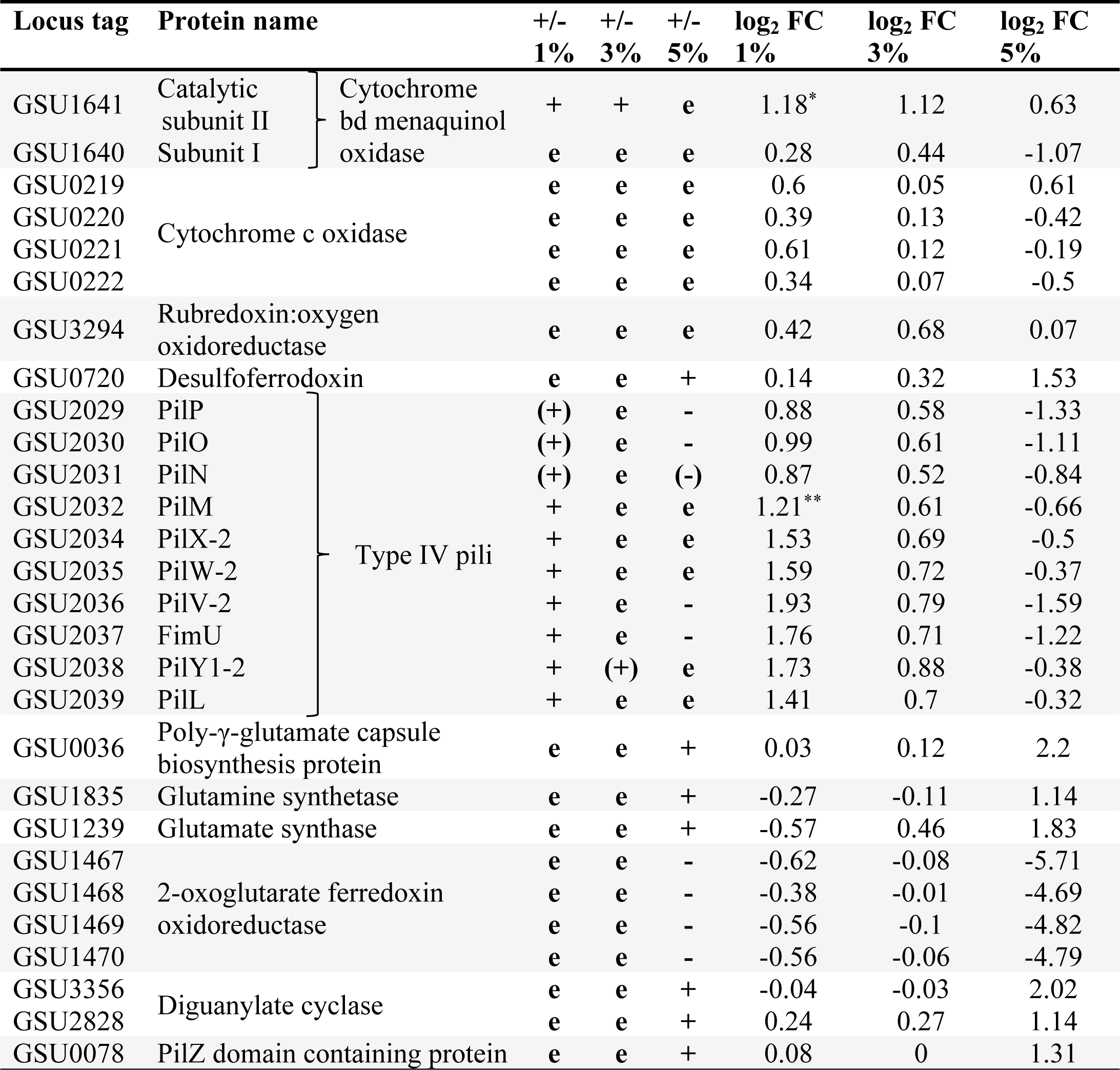
Selected genes expressed differently under microaerobic conditions. Differentially expressed genes with p-value >0.05 and >0.1 are marked * and ** respectively. 1%, 3% and 5% refer to experiments 4F1%, 4F3% and 4F5% respectively, + = higher expressen, - = lower expression, e = not differentially expressed, FC = fold change.

## Discussion

### Comparison of microaerobic and anaerobic growth

*G. sulfurreducens* was cultivated under microaerobic and anaerobic conditions in a bioreactor system. Cell growth could be observed in both the strictly anaerobic as well as the microaerobic cultivations. Under electron acceptor limiting conditions (experiments 4F0%, 4F1% and 4F3%), cell growth deviated from exponential growth and behaved linear. The increase in cell dry weight (CDW) occurred faster when oxygen was provided in comparison to the anaerobic experiment 4F0%. Also, when comparing experiments 4F1% and 4F3% directly, it becomes apparent that a higher oxygen feed also resulted in faster growth of the bacteria. The maximum biomass concentration achieved in experiment 4F3% was the same as in the anaerobic control experiment 40F0%. Thus, oxygen is clearly a suitable electron acceptor for *G. sulfurreducens,* enabling cell growth on acetate to the same extent as with fumarate.

In all successful microaerobic cultivations, the DO concentration was around 0 mg L^-1^ during the time of cell growth and acetate consumption. There was only cysteine as oxygen capturing chemical species present in the cultivation medium, but due to the overnight equilibration of reactors at aerobic conditions, cysteine should already be completely reduced. This suggests that *G. sulfurreducens* used oxygen as terminal electron acceptor. As oxygen was fed continuously at the same concentration and was consumed completely, this compound was limiting for cell growth, which explains the resulting linear increase of the biomass concentration and the respective linear decrease of acetate.

The rise in DO after about 24 h in experiment 4F3% was in accordance with the stop of bacterial growth and the ceasing acetate consumption. After acetate was depleted, cell growth could no longer be sustained due to lack of another carbon and energy source. Likewise, no electrons were available to be transferred to oxygen. The maximum DO concentration when gassing with 3 % of oxygen was determined to be 0.93 mg L^-1^. The actually measured DO concentration after cell growth subsided was approximately 1 mg L^-1^ and was thus quite close to the expected maximum. Thus, as a result of acetate depletion, the entire amount of oxygen remained solved in the cultivation medium.

When 5 % of oxygen was fed in the gas inlet no cell growth was observed and the DO concentration rose to the expected maximum DO, indicating that there was no, not even a partial, oxygen reduction. This suggests there is a limit of oxygen manageable by the cells, which must be in the range between 3 to 5 % of oxygen in the gas inlet. When this limit is exceeded, oxygen is not consumed by the cells at all and cell growth is inhibited entirely. The growth with oxygen can thus be described as an “all-or-nothing” state for *G. sulfurreducens*.

The use of oxygen as terminal electron acceptor by *G. sulfurreducens* was already described previously by Lin et al., who observed the consumption of oxygen and acetate in cultivations initially grown anaerobically with fumarate as electron acceptor (2). They observed a similar dependency of the ability of oxygen reduction by *G. sulfurreducens* on the provided oxygen concentration. Consumption of oxygen and cell growth were only achieved when 5 or 10 % of oxygen were added to the headspace of culture flasks, but not at higher concentrations (2). These authors only measured oxygen in the head space of culture flasks and did not provide information on DO concentrations. Thus, only a qualitative comparison of Lin et al. with the results presented in this study is possible.

The capability of *G. sulfurreducens* to deplete oxygen is likely dependent on the amount of cells present. Therefore, an experiment was conducted in which the oxygen concentration was continually raised depending on the biomass concentration. The maximum sOUR was calculated as described in the materials and methods section to be 95 mg_O2_ g_CDW_ ^-1^ h^-1^. The sOUR was kept at this level throughout experiment 4FSI by a stepwise increase of the gas inlet oxygen concentration dependent on the observed CDW at each time of sampling. In this way, a final gas inlet concentration of 12.5 % of oxygen was achieved, which is 2.5 times higher than the here beforehand determined growth inhibiting oxygen concentration of 5 % in the gas inlet. This clearly demonstrates that the amount of oxygen usable by *G. sulfurreducens* is dependent on the amount of biomass. This result is novel to the study of Lin et al., who just showed a continuous cell growth on oxygen and acetate, when 5 % of oxygen was repeatedly provided in the head space of cultivations (2).

The maximum specific growth rate achieved in experiment 4FSI was determined to be 0.124 ± 0.007 h^-1^, which is in the same range as the specific growth rate determined for the anaerobic conditions with excess of fumarate to be 0.109 ± 0.015 h^-1^. This result further illustrates the suitability of oxygen as a terminal electron acceptor for *G. sulfurreducens*. The achieved maximum CDW after 20 h of cultivation is 13 % higher in this experiment than in the anaerobic control experiment (40F0%). Therefore, oxygen seems to affect the cells in a way that allows for a more efficient substrate utilisation, which will be further discussed in the next section.

### Concentration profiles of selected organic acids and implications for metabolism

The concentrations of acetate, succinate, fumarate and pyruvate were determined during cultivations. Their concentration profiles can give insight into the metabolism of *G. sulfurreducens* under anaerobic and microaerobic conditions.

Acetate was consumed in all cultivations for biomass and energy production. The only mechanism for ATP generation in *G. sulfurreducens* is by electron transport phosphorylation (9,10). Hereby, the greater potential energy arising from the high redox potential of oxygen (815 mV) (11) compared to the redox couple fumarate/succinate (30 mV) (12) could allow for a more efficient acetate utilisation of *G. sulfurreducens* when grown under microaerobic conditions. To investigate this further, the biomass yield coefficient for acetate as substrate (*Y*_*X/S*_) was calculated from the linear slope of the biomass concentration plotted against the acetate concentration during the growth phase of each experiment. *Y*_*X/S*_ was at 0.552 ± 0.014 g_CDW_ g_s_ ^-1^ under anaerobic conditions (40F0%) and was similar for all cultivations except 4F1%, which yielded a slightly higher value of 0.63 ± 0.06 g_CDW_ g_s_ ^-1^. This indicates that the use of oxygen as electron acceptor might indeed enable the cells to utilise acetate more efficiently, however only if the sOUR is low. Since no cell growth and oxygen reduction could be observed in experiment 4F5%, not even a partial reduction, the priority for *G. sulfurreducens* when confronted with oxygen will apparently be to annihilate all oxygen, independent of the potential energetic benefits. This might explain why *Y*_*X/S*_ returns to 0.55 g_CDW_ g_s_ ^-1^, when higher concentrations of oxygen are provided. Similarly, Lin et al. also determined the same biomass yield per mole of acetate with either fumarate or oxygen as electron acceptor (2). It may be that the amount of oxygen in the headspace of 5 % or 10 % provided by these authors (2) was already too high to allow for an investigation of the increased *Y*_*X/S*_ that was observed in this study for cultivation 4F1%.

The yield of oxygen reduced per mole of acetate *Y*_*O2/S*_ was determined to be 1.76 ± 0.12 mol_O2_ mol_s_ ^-1^ which is in the same range as, but more precise than the value of 2.4 ± 0.8 mol mol^-1^ determined by Lin et al. (2). If all acetate is degraded to CO_2_, *Y*_*O2/S*_ should be 2 mol_O2_ mol_s_ ^-1^. If a complete degradation of acetate to CO_2_ was taking place, however, no carbon would be available for the formation of biomass and of other substrates. Thus, the value determined here is quite reasonable.

For all cultivations, a consumption of fumarate and a simultaneous production of succinate can be observed. Under anaerobic conditions, fumarate is used as terminal electron acceptor and reduced to succinate via a bifunctional fumarate reductase / succinate dehydrogenase (12). The tricarboxylic acid (TCA) cycle is functioning as an open loop under these conditions (see **Fig. 5**, black lines) (10,12,13). The stoichiometry of this reaction is 1:1 and accordingly a molar succinate (Suc) to fumarate (Fum) yield (*Y*_*Suc/Fum*_) of 0.972 ± 0.025 mol_Suc_ mol_Fum_ ^-1^ is seen in cultivation 40F0%. With only 4 mM of fumarate supplemented, however, *Y*_*Suc/Fum*_ increases to 1.30 ± 0.05 mol_Suc_ mol_Fum_ ^-1^. Thus, more succinate was formed here than it should have been possible from the provided fumarate. This indicates that other substrates were used by the cells as precursor for fumarate, to create more of the electron acceptor. The declining pyruvate concentration in this experiment, for example, indicates an enhancement of anaplerotic reactions, which would ultimately lead to fumarate (see **Fig. 5**, red lines). However, it should be noted that the amount of pyruvate found in the medium is not sufficient to account for all observed succinate, so it should be considered that a different carbon source is responsible for the succinate accumulation. This could be cysteine provided in the medium, as cysteine can be degraded to pyruvate by *G. sulfurreducens* (3), thus providing an additional carbon source for fumarate.

**Fig. 5.**
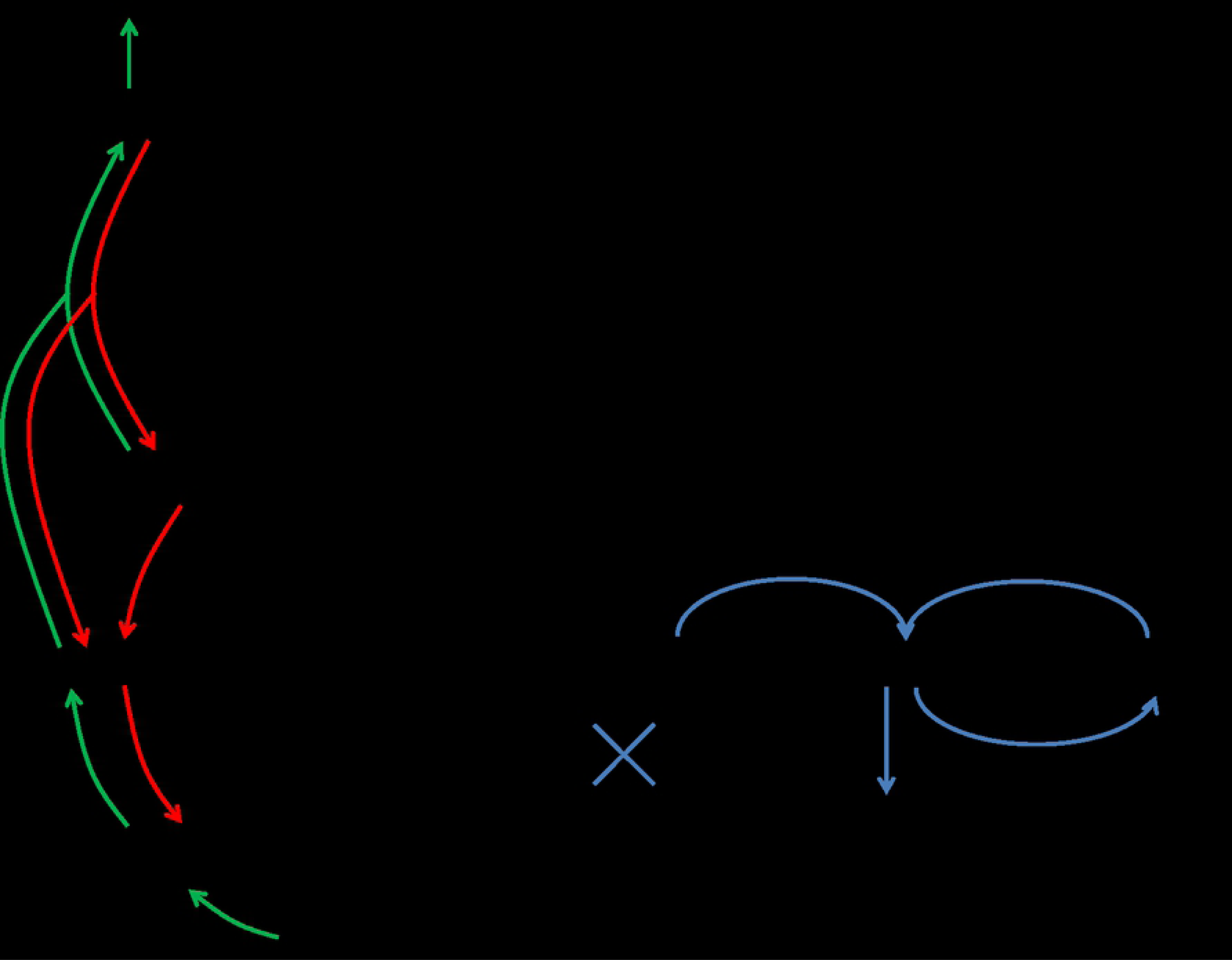
Model of changes in carbon flow during growth of *G. sulfurreducens* under electron acceptor limiting (red), oxygen reducing (green) and oxygen tolerating (blue) conditions compared to anaerobic growth with excess of fumarate (black).

With the addition of oxygen, the production of succinate from fumarate turns into a consumption. This consumption increases with increasing oxygen concentration as seen in **Fig. 4**. As oxygen now serves as electron acceptor, the bifunctional fumarate reductase / succinate dehydrogenase can operate in the other direction and the TCA cycle can function as a closed loop (see **Fig. 5**, green lines).

The concentration profiles of pyruvate support the beforehand discussed implications for the TCA cycle. There is an increase of pyruvate observed when the sOUR is high. This corresponds well to the degradation of succinate via the TCA cycle. The complete reduction of oxygen observed together with the fact that cell growth is absolutely inhibited when this reduction does not take place indicates a need of *G. sulfurreducens* to dispose of any oxygen found in the surrounding of the cells. Thus, with a high sOUR, the flow of carbon in the TCA cycle is shifted from succinate towards oxaloacetate and pyruvate, whereby two more redox equivalents (menaquinol and NAD(P)H) are produced (12). Those may in turn serve as electron donor to reduce oxygen to water.

Pyruvate derived from succinate can serve as an additional carbon source for cell growth. During the first 18 h of the cultivation with stepwise increasing oxygen concentrations, 3 mM of succinate are being consumed, but only about 0.5 mM of pyruvate are formed. Assuming a 1:1 stoichiometry for this transformation, there are potentially up to 2.5 mM of pyruvate available for further biomass formation, which can explain the 13 % increased cell dry weight found in this cultivation that was observed even though the biomass/acetate yield did not change significantly with oxygen as terminal electron acceptor (see **Fig. 5**, green lines). To fully account for all carbon, however, a more precise analysis of intracellular metabolites and their fluxes as well as the determination of the elementary composition of *G. sulfurreducens* is necessary.

### Transcriptome analysis

Transcriptome analysis was performed to determine the mechanism by which *G. sulfurreducens* reduces oxygen and to estimate strategies for survival of *G. sulfurreducens* when exposed to oxygen.

Several genes have been found in the genome of *G. sulfurreducens* encoding for enzymes that may enable this organism to reduce oxygen (3). Usually in aerobic bacteria, cytochrome bc1 complex and cytochrome c oxidase, which represent complexes III and IV of the oxidative phosphorylation pathway, are responsible for the transportation of electrons from the membrane bound quinon pool to a cytochrome c, which then functions as electron donor for the reduction of oxygen to water. The cytochrome c oxidase was formerly suspected to be the enzyme responsible for oxygen reduction in *G. sulfurreducens* (2). Núñez et al. showed that the gene for this enzyme is part of the RpoS regulon which is necessary for growth of *G. sulfurreducens* with oxygen (5). These authors even reported that cell growth with oxygen was impossible without cytochrome c oxidase (W. Lin, unpublished results according to (5)). The corresponding genes were, however, not expressed at higher levels under microaerobic conditions in the microarray analysis performed here. Another oxygen reducing enzyme whose gene is present in the *G. sulfurreducens* genome and is also regulated by RpoS is a putative rubredoxin:oxyen oxidoreductase (3,5). However, the corresponding gene was not differentially expressed under any of the conditions investigated here.

Instead, transcriptome analysis revealed the up to 2.3-fold upregulation of the expression of the gene GSU1641 in experiments 4F3% and 4F1% which encodes the catalytic subunit II of the cytochrome bd complex menaquinol oxidase, an enzyme responsible for the reduction of O_2_ to H_2_O and the simultaneous oxidation of menaquinol to menaquinon. The expression of GSU1641 was also found to be regulated by RpoS by Núñez et al. (there named cytochrome d ubiquinol oxidase) (5). In another study that investigated *G. uraniireducens*, there was also a menaquinol oxidase among the genes expressed at higher levels when *G. uraniireducens* was exposed to oxygen (6). This suggests the menaquinol oxidase encoded by GSU1641 to be the enzyme most important for oxygen reduction in *G. sulfurreducens* rather than the conventional oxidative phosphorylation pathway. It is possible that the other potential oxygen reducing enzymes may also be involved in oxygen reduction by *G. sulfurreducens*. But to reduce oxygen entirely, *G. sulfurreducens* relies on a higher expression of only the gene for the menaquinol oxidase. The reason for this may be that the menaquinol oxidase is a high-oxygen affinity terminal oxidase (5), thus ensuring a fast and efficient removal of any intruding oxygen.

Several other genes found in the RpoS regulon were not differentially expressed in this study except for a gene encoding for desulfoferrodoxin (GSU0720) which was expressed at 2.9-fold higher levels in experiment 4F5%. Desulfoferrodoxins are enzymes that reduce superoxide to hydrogen peroxide and are often found in anaerobic bacteria in association with protection against oxidative stress (14,15). Thus, the higher expression of this gene is in accordance with the lack of cell growth and oxygen reduction in this experiment.

Genes for other components required for energy metabolism like the ATP synthase (GSU0108-GSU0114) or the NADH dehydrogenase (GSU0338-GSU0351) were not expressed differentially in experiment 4F3%, suggesting an otherwise unimpaired energy production machinery (see supplementary material). Likewise, there were no differences with genes for enzymes of the TCA cycle or the glycolysis/gluconeogenesis pathway, indicating that growth and metabolism of *G. sulfurreducens* work in principle in the same way as under anaerobic conditions. This result corresponds well to the observed similar cell growth at 3 % of oxygen and also in experiment 4FSI.

Contrary to this, the expression of many genes of the oxidative phosphorylation, the TCA cycle and the glycolysis/gluconeogenesis is downregulated at 5 % of oxygen, corresponding to an inhibition of cell growth seen in this experiment. For example, the expression of GSU0338 to GSU0351 encoding for the NADH dehydrogenase was downregulated up to 23-fold and GSU1466 which encodes for malate dehydrogenase was expressed at 28-fold lower levels (see supplementary material). On the other hand, expression of genes associated with oxidative stress (e.g. the genes for the ferritin-like domain containing proteins, GSU2193 and GSU2967, and for a universal stress protein, GSU0515) were upregulated up to 9-fold.

Transcriptomic analysis also yielded results that give insight into strategies for survival under oxygen exposure depending on the oxygen concentration. Experiment 4F1% resulted in the up to 3.8-fold upregulation of the expression of 11 type IV pilus genes (GSU2029-GSU2039). Type IV pili have multiple functions in bacteria, including cell communication, adhesion, biofilm formation and cell motility (16,17). Motility is caused by cells protruding pilus fibres whose tips attach on a surface and subsequently retracting the pilus again, pulling the cell body forward (18). In *G. sulfurreducens*, pili have another special function of serving as conductive wires for electron transport to extracellular acceptors (19). As both the electron acceptor fumarate and oxygen were rather limited in experiment 4F1%, the production of pili could have been a means of cells to find other, insoluble electron acceptors. Insoluble iron oxides are hypothesised to bind to the pili and to be reduced by electrons that are transferred by the conductive major pilin PilA (20). The reduced iron oxides are then shed off by retraction of the pilus, which is powered by the ATPase PilT-4 (20). The genes for these two proteins together with other type IV pilus genes are, however, part of a different cluster located elsewhere in the genome (GSU1491-GSU1497) and are not part of the genes found expressed at higher levels here.

While the above mentioned type IV pili genes have been investigated by several additional studies (21–23), the genes encoded by GSU2029 to GSU2039 have to our knowledge not been investigated more closely in *Geobacter* species. GSU2029 to GSU2039 include genes for PilM, PilN, PilO and PilP, which are thought to be part of the assembly responsible for pilus protrusion and retraction in *G. sulfurreducens* (19) as they serve this function in other bacteria (16,18). Some of the remaining genes encode for PilX-2, PilW-2, PilV-2 and FimU, which represent minor pilins. In *Pseudomonas aeruginosa*, minor pilins are thought to form initiation complexes from which the protrusion of a new pilus can commence (18). Displaying more minor pilins in the membrane would result in more, but shorter, pili protruding from the cells (18). More pili could enable *G. sulfurreducens* to attach to a surface more efficiently in order to use the above described retraction mechanism for motility. Thus, they could be used to flee the microaerobic environment. Expression of these genes returned to the levels found under anaerobic conditions with the increase of oxygen to 3 %. As protrusion and retraction of pili is an ATP intensive process, *G. sulfurreducens* might only follow the strategy of fleeing the zone of oxygen, when the concentration is adequately low and fleeing may actually result in finding an anaerobic zone. Further analysis is needed to assess the exact role of these pili proteins in *G. sulfurreducens*, especially since *G. sulfurreducens* was originally described to be non-motile (1).

In experiment 4F5%, transcriptome analysis revealed higher expression of genes encoding for enzymes responsible for biofilm formation and encapsulation. For example, GSU0036 encodes for a poly-γ-glutamate capsule biosynthesis protein and was expressed at 4.6-fold higher levels. Correspondingly, the expression of genes encoding for enzymes involved in glutamate metabolism was likewise upregulated up to 3.6-fold, i.e. the gene for glutamine synthetase (GSU1835) and for the glutamate synthase (GSU1239). Some of the genes whose expression was most strongly downregulated (up to 52-fold) in experiment 4F5% are encoding for a 2-oxoglutarate ferredoxin oxidoreductase (GSU1467-1470) indicating an inhibition of the carbon flux from 2-oxoglutarate to succinyl-CoA to allow for the formation of glutamate (see **Fig. 5**, blue lines).

Another indicator for biofilm formation is the higher expression (up to 4-fold) of GSU0078, which encodes for a PilZ domain containing protein and GSU3356 as well as GSU2828, which both encode for diguanylate cyclase, an enzyme producing cyclic diguanylate (c-di-GMP). C-di-GMP has been reported to function as signalling molecule when binding to the PilZ-domain of bacterial pili, inhibiting movement and inducing biofilm formation (24–26). Another indicator for inhibition of motility is the 2.2-fold upregulation of GSU2185, a negative regulator for the production of flagellin (see supplementary material). Further, the type IV pili genes whose expression was found to be upregulated in experiment 4F1% were mainly expressed at lower levels in 4F5% (see **Table 1**). These results suggest that *G. sulfurreducens* follows the strategy of encapsulation rather than fleeing when confronted with concentrations too high to allow for oxygen reduction. These results correspond well with results from Mouser et al., who found genes involved in motility to be expressed at lower levels in *G. uraniireducens* when exposed to 5 % of oxygen (6).

## Conclusion

The data presented here show that *G. sulfurreducens* can grow with oxygen as the terminal electron acceptor with a maximum sOUR of 95 mg_O2_ g_CDW_ ^-1^ h^-1^. The cytochrome bd complex menaquinol oxidase was found to be the enzyme most likely responsible for the main oxygen reduction. Thus, the bacterium should be classified as facultative microaerobe rather than strict anaerobe. With oxygen as terminal electron acceptor, the TCA cycle functions as a closed loop, enabling *G. sulfurreducens* to utilise succinate as a carbon source for biomass production. Transcriptome analysis has revealed a tendency of *G. sulfurreducens* to escape low concentrations of oxygen and to react to higher oxygen concentrations by encapsulation or biofilm formation. The results presented here fit well into the already existing studies dealing with the reaction of *Geobacter* spp. to oxidative stress or oxygen reduction and help to further understand these organisms.

## Materials and Methods

### Bacterial strain and cultivation medium

*Geobacter sulfurreducens* PCA (DSMZ 12127) was used in this study and was obtained from the German Collection of Microorganisms and Cell Cultures (DSMZ), Braunschweig, Germany.

*G. sulfurreducens* was cultivated in a minimal medium containing 10 mM acetate as carbon source and electron donor and 40 mM fumarate as electron acceptor as described previously (27–29). For precultures medium was prepared in serum bottles (125 mL with 50 mL liquid volume, Fisher Scientific, Hampton, New Hampshire, USA) and gassed with N_2_ for a minimum of 20 min to obtain anaerobic conditions. Subsequently, 20 % (v/v) of a consisting *G. sulfurreducens* culture was added as inoculum and serum bottles were sealed with butyl rubber stoppers. Cells were incubated at 180 min^-1^ (25 mm deflection, Multitron, Infors AG, Bottmingen, Switzerland) and 30 °C for about 24 h.

### Microaerobic cultivation in bioreactor setup

For microaerobic cultivations, 1 L bioreactor systems (DASGIP Information and Process Technology GmbH, Jülich, Germany) with a working volume of 500 mL were used. Medium was prepared as described above with altering fumarate concentrations (see **Table 2**) and sterilised at 121 °C and 2 bar (200,000 Pa) for 20 min. After sterilisation, vitamin solution, trace mineral solution, and L-cysteine were added in a sterile manner. Bioreactors were set to operating conditions (Rushton turbines at 200 min^-1^, temperature of 30 °C, gassing with air (21 % (v/v) oxygen) at a gasing rate of 6 sL h^-1^) overnight to allow for equilibration of the DO sensor (Visiferm DO 225, Hamilton, Höchst im Odenwald, Germany) before calibration. DO sensors were calibrated by a two-point calibration using 21 % oxygen and 0 % oxygen (corresponding to 100 % N_2_), respectively, in the gas inlet. Before inoculation, anaerobic conditions were ensured by continuous gassing with N_2_ and verified with the DO sensor signal. As inoculum, 100 mL of a *G. sulfurreducens* preculture were added sterilely.

**Table 2.** Experiments conducted for comparison of anaerobic and microaerobic growth of *G. sulfurreducens*.

| Experiment abbreviation | Fumarate [mM] | Oxygen in gas inlet [%] |
| --- | --- | --- |
| 40F0% | 40 | 0 |
| 4F0% | 4 | 0 |
| 4F1% | 4 | 1 |
| 4F3% | 4 | 3 |
| 4F5% | 4 | 5 |
| 4FSI | 4 | stepwise increase from 3 to 12.5 |

Cultivations were started with an initial anaerobic period of 2 h. For microaerobic cultivations, the gas composition was changed after this 2 h period to contain oxygen according to **Table 2** by mixing of N_2_ and air. Biomass concentration was determined by periodic sampling (1 mL, 2 h interval) and measurement of the optical density (Libra S11, Biochrom, Berlin, Germany) at 600 nm (OD_600_). For conversion into cell dry weight (CDW) a beforehand determined conversion factor of 0.62 g_CDW_ L^-1^ OD_600_ ^-1^ was used. In one experiment, the gas inlet concentration was changed after OD_600_ measurement depending on the determined CDW to maintain a specific oxygen uptake rate (sOUR) of 95 mg_O2_ g_CDW_ ^-1^ h^-1^ (experiment abbreviation: 4FSI). Experiments were performed in biological triplicates.

### Determination of maximum dissolved oxygen concentration, volumetric oxygen transfer coefficient and maximum specific oxygen uptake rate

To allow for quantification of oxygen, the volumetric oxygen transfer coefficient (*k*_*L*_ *a*), and the maximum dissolved oxygen (DO) concentration *c*^*\**^ _*O2*_ were determined for the bioreactor system and medium used. *c*^*\**^ _*O2*_ was approximated mathematically to be 0.31 mg L^-1^ per % of oxygen in the gas inlet at 30 °C with a method described previously (30). *k*_*L*_ *a* was determined using cultivation medium without cells with the dynamic gassing out method, whereby oxygen was stripped from the medium with nitrogen and subsequently the response of the DO sensor was monitored during gassing with oxygen concentrations of 3, 5 and 21 % (air). From the linear slope of *ln(c*^*\**^ _*O2*_ *– c*_*O2*_ ) plotted against the time the *k*_*L*_ *a* was calculated to be 9.4 ± 1.1 h^-1^.

Provided that complete oxygen reduction is occurring, the specific oxygen uptake rate (sOUR) can be determined by the following equation 1 from the identified *kla*-value, the determined maximum DO concentration *c*^*\**^ _*O2*_ and the observed cell dry weight *X*.

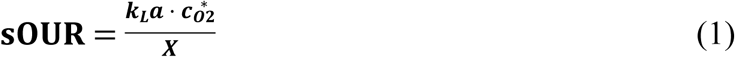

The maximum sOUR was determined using equation **Error! Reference source not found.** and data obtained in experiment 4F3% at 2 h of cultivation (*c*^*\**^ _*O2*_ of 0.935 mg_O2_ L^-1^ and *X* of 0.0928 ± 0.0024 g_CDW_ L^-1^) to be 95 ± 11 mg_O2_ g_CDW_ ^-1^ h^-1^. This experiment and time-point were chosen as the highest sOUR was observed here.

### Determination of organic acid concentration

Acetate, fumarate, succinate, pyruvate and 2-oxoglutarate were determined using an HPLC system (LaChrom Elite®, Hitachi, Tokyo, Japan) equipped with an Aminex® HPX-87H column (Bio-Rad Laboratories, Hercules, California, USA) as the stationary phase and 12.5 mM H_2_ SO_4_ as mobile phase at 0.3 mL min^-1^ and 25 °C at retention times of approx. 33, 35, 26, 21 and 20 min, respectively. Detection was performed using a refractive index detector and a diode array detector at 210 nm or 230 nm (only for fumarate). Final concentrations were calculated as the mean of the signal from both detectors.

### Analysis of transcriptome

Samples for RNA analysis were acquired during cultivations by centrifugation of 15 mL or 50 mL of cultivation broth at 11,800 g and 4 °C for 10 min. After discarding the supernatant, cell pellets were immediately frozen in liquid nitrogen and stored at −80 °C until RNA purification.

Cells were disrupted enzymatically and mechanically using a similar protocol as described previously (31). Precipitated cells were resuspended in 200 µL of TE-buffer (10 mM Tris HCl, 1 mM EDTA, pH 8) containing lysozyme from chicken egg white (Sigma Aldrich, St. Louis, Missouri, USA) at a concentration of 15 mg mL^-1^ and incubated for 30 min at ambient temperature with regular mixing. Subsequently, glass beads (diameter 150 – 212 µm) were added and cells were vortexed for 3 min for mechanical cell disruption. RNA was purified from the supernatant using the RNeasy Mini Kit (Quiagen, Hilden, Germany) according to the manufacturer’s instructions. Digestion of chromosomal DNA with DNAse (Quiagen, Hilden, Germany) was carried out twice. RNA concentration was determined using the NanoDrop ND-1000 (Peqlab Biotechnology, Erlangen, Germany) and RNA quality was assessed with the 2100 Bioanalyzer (Agilent Technologies, Santa Clara, California, USA) according to the manufacturer’s instructions. Samples with an RNA integration number of 7 or higher were used for transcriptome analysis.

Purified RNA was stained with cyanine 3 or cyanine 5 using the ULS Fluorescent Labeling Kit for Agilent arrays from Kreatech (Leica Biosystems, Amsterdam, Netherlands) according to the manufacturer’s instructions. The degree of labelling was determined using the NanoDrop ND-1000 (Peqlab Biotechnology, Erlangen, Germany) and RNA with a degree of labelling between 1 and 3.6 was used for analysis. Subsequently, RNA was fragmented and hybridized to a custom-made GE 8×15K microarray for analysis of gene expression of *G. sulfurreducens* (Agilent Technologies, Santa Clara, California, USA) using the gene expression hybridization kit from Agilent (Agilent Technologies, Santa Clara, California, USA) according to the manufacturer’s instructions. Subsequently, the microarray was analysed using the Microarray C Scanner (Agilent Technologies, Santa Clara, California, USA) as described previously (32). Genes with a |logFC| of >1.0 were considered differentially expressed. All data have been deposited in the NCBI Gene Expression Omnibus (GEO). The GEO Series accession number is GSE124792. Reviewers can use the following token to get access: **qtylsgmatxetbib**.

## Acknowledgements

Special thanks go to Annika Michel of the Institute of Microbiology at the Technische Universität Braunschweig for help with performing RNA purification and microarray preparation.

## Supporting Information

**S1_Fig. Concentration profiles of pyruvate during growth of *G. sulfurreducens* at 30 °C (A) during experiments 4F0%, 4F1%, 4F3% and 4FSI (4 mM of fumarate and altering oxygen concentrations of 0 %, 1 %, 3 % and stepwise increasing oxygen concentrations, respectively) as well as (B) during experiment 40F0% (40 mM of fumarate and 0 % of oxygen). Error bars indicate standard deviation of biological triplicates.**

with altering oxygen (between 0 and 5 %) and fumarate (4 or 40 mM) concentrations

**S1_Table. Transcriptome data of *G. sulfurreducens* grown under microaerobic conditions with 4 mM of fumarate and 1 % of oxygen provided in the gas inlet as electron acceptors. Data were compared to *G. sulfurreducens* grown under anaerobic conditions with 40 mM of fumarate as electron acceptor.**

**S2_Table. Transcriptome data of *G. sulfurreducens* grown under microaerobic conditions with 4 mM of fumarate and 3 % of oxygen provided in the gas inlet as electron acceptors. Data were compared to *G. sulfurreducens* grown under anaerobic conditions with 40 mM of fumarate as electron acceptor.**

**S3_Table. Transcriptome data of *G. sulfurreducens* grown under microaerobic conditions with 4 mM of fumarate and 5 % of oxygen provided in the gas inlet as electron acceptors. Data were compared to *G. sulfurreducens* grown under anaerobic conditions with 40 mM of fumarate as electron acceptor.**

